# Mammalian Cell Models of the Distal Convoluted Tubule: A Systematic Review of Cell Lines, Culture Condition and Gene Expression

**DOI:** 10.1101/2024.02.23.581724

**Authors:** Chutong Zhong, Zhen Sun, Alessandra Grillo, Stephen B Walsh, Keith Siew

**Affiliations:** London Tubular Centre, Department of Renal Medicine, University College London, London, UK

**Author notes:** Correspondence: *Keith Siew (*).

**Keywords:** distal convoluted tubule, cell models, renal physiology, ion transport and channels, hypertension

## Abstract

This review examines the crucial role of human cellular models in renal physiology research, with a specific focus on the distal convoluted tubule (DCT). It aims to provide a comprehensive summary of the origins, culture practices, and genetic studies associated with commonly employed DCT cell models. To achieve this, a systematic literature review was performed on Europe PMC, employing a Boolean search strategy. A total of 6,559 articles were initially screened, resulting in 301 articles on human-origin cell models, 69 on murine models, and 29 on canine models being included in the final analysis. Notably, the review identified two studies that introduced novel immortalised human DCT cell lines developed from primary DCT cells. This paper provides a detailed account of the lineage of each cell model, their prevalent culture conditions, and the frequency and nature of gene transfections—both wild type and mutant—conducted in these models. A significant observation from the analysis was the inconsistent reporting of methodological details across studies, which compromises the reproducibility of the research. Additionally, there was a considerable variation in culture conditions and transfection methods used across different studies. The review also highlights that HEK293 family and murine DCT cell lines do not serve as accurate models for DCT cells, pointing out the frequent need for transfecting DCT-specific genes to simulate DCT functionality adequately.

**NEW & NOTEWORTHY:** This review is the first review to detail the current state of play with cell models of the distal convoluted tubule, including endogenous protein expression, culture conditions etc. We expect this review to be of great utility to established and early career researchers who are interested in nephrological or hypertensive patho/physiology. We have highlighted some important gaps in reporting the established cell models, we think that this kind of review is important for open science.

## INTRODUCTION

Hypertension, a significant etiological factor implicated in a multitude of cardiovascular and renal pathologies (1), is expected to escalate by 60%, with the number of people affected surpassing 1.5 billion in the next two decades (2). Recent research on Gordon and Gitelman syndrome, two uncommon kidney disorders marked by opposite effects on blood potassium levels and blood pressure due to over- and under-activation respectively of the thiazide-sensitive sodium-chloride transporter (*SLC12A3,* NCC), has highlighted the role of the distal convoluted tubule (DCT) in controlling blood pressure (3–7). The with-no-lysine (WNK) kinases, WNK1 and WNK4, STE20-related proline alanine-rich kinase (SPAK) and oxidative stress response 1 kinase (OxSR1) are expressed in the DCT and regulate the activity of NCC through phosphorylation of key residues to control Na reabsorption and consequently blood pressure (8, 9).

Several mammalian renal tubular cell lines (human, murine and canine) have been used to explore the interplay between electrolyte homeostasis and blood pressure. Human Embryonic Kidney 293 (HEK293), Murine Distal Convoluted Tubule 209 (mDCT209), Murine Distal Convoluted Tubule 15 (mDCT15), Transgenic Mouse (SV-PK/Tag) Distal Convoluted Tubule Cells (mpkDCT), Madin-Darby Canine Kidney (MDCK) and their derivatives are most frequently utilised to model human DCT physiology *in vitro.* Primary culture, although less common and more challenging, is another favored means of studying DCT function. Theoretically, these primary cell *in vitro* models should better reflect *in vivo* DCT physiology, however significant variations in their means of isolation and culture condition from lab to lab, as well as inconsistencies in endogenous gene expression can make interpretation of results and subsequent replication of findings challenging (10).

The purpose of this systematic review is to summarise and compare the culture conditions, phenotypic characteristics and commonly studied gene products of extant DCT cell models published up until the year 2021. This resource will enable researchers to make informed decisions about their choice of experimental models and to contextualise the findings of past and future experiments.

## MATERIALS AND METHODS

A literature search of English-language papers was conducted using Europe PMC for all the original research papers published up until 24 June 2021. The search strategy involved using Boolean search functions “AND/OR” to combine and integrate into a single search string: the full name, abbreviations and synonyms for the cell lines, DCT related terms and wildcards, canonical DCT-specific genes and gene products. For HEK293 and related derivatives, the keywords of cell lines included HEK293*, HEK293T and HEK293-TREx. For murine and canis cell lines, the keywords would be mDCT209, mDCT15, mpkDCT and MDCK*. All keywords of cell lines were combinedly searched with WNK*, SPAK*, TSC, NCC, NKCC*, OSR1, KLHL3, MO25, ROMK*, CAB39, SLC12A3, Cul3, Kir4.1/5.1. The search string iteratively refined using the Boolean function “NOT” to exclude irrelevant expansions for some abbreviations (e.g. OSR1 may also refer to ‘odd-skipped related transcription factor 1’). Original research papers exclusively performing *in vivo* human or animal model experiments, *ex vivo* intact tissues or those encompassing research with little relevance to the study of renal function, blood pressure or electrolyte disorders were excluded, in addition to review articles and conference abstracts. A schematic representation of paper selection is detailed in **Figure 1**. Further manual data validation was performed by three people to ensure compliance to the inclusion criteria and removal of duplicates.

**Figure 1.**
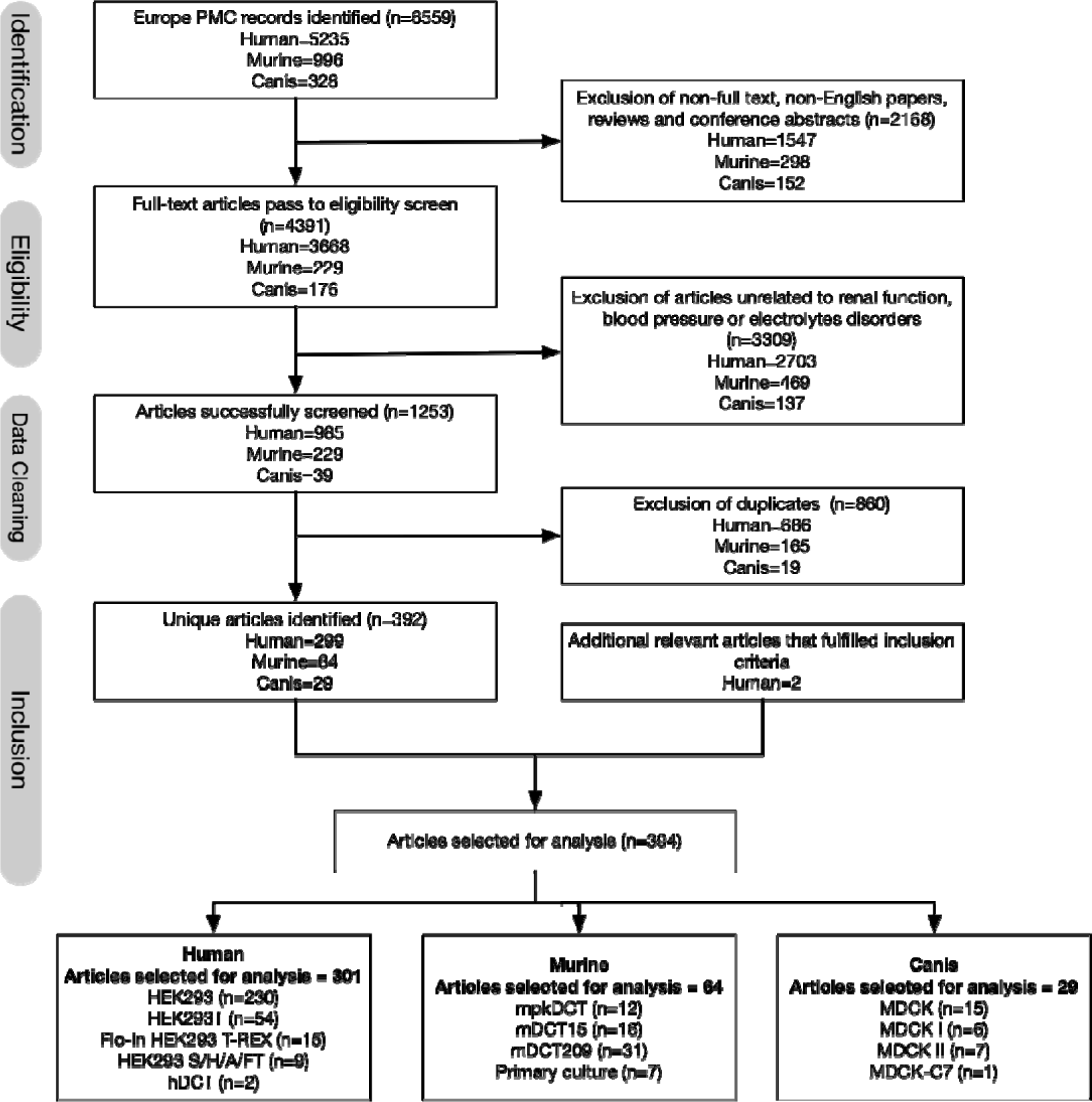
Algorithm for search and selection of articles using EuropePMC. hDCT=human DCT cell model. Two relevant papers that were known to the authors but not detected by Europe PMC were additionally included.

Culture conditions, transfection methods and genes of interest were extracted from the original papers where available. Terminology, scientific units and gene names were standardised for compiling of the descriptive statistics. Duplicated culture conditions in each cell line were summarised and ranked by popularity.

## RESULTS

The literature search resulted in 6559 initial hits. A total of 394 papers met the inclusion criteria, comprising of 301 papers for human DCT cell models, 64 papers for murine and 29 papers for canine. Only two papers were identified using immortalised human DCT cells (11, 12). A detailed breakdown of the species distribution across cell lines reported in the literature are presented in **Figure 2**.

**Figure 2.**
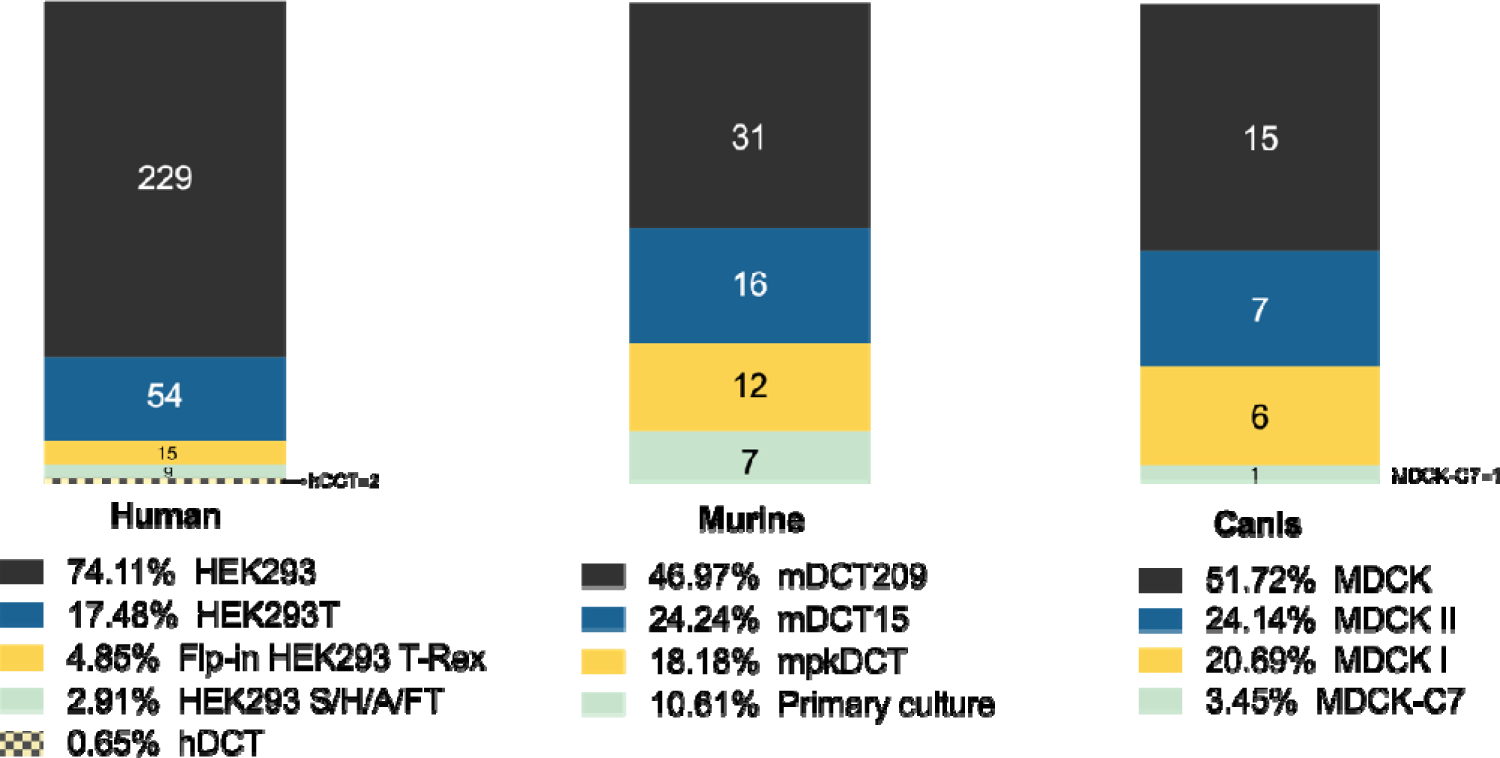
Overview of number of cell lines identified in selected papers grouped by species. **Human:** Total conditions included for analysis using human DCT cell lines=309. **Murine**: Total conditions included for analysis using murine DCT cell lines=66. **Canis**: Total conditions included for analysis using canine DCT cell lines=29.

### Culture Condition: Human Cell Lines

Culture conditions were sorted into culture medium, serum, antibiotics, supplements and gas mixture (**Figure 3**). Any category for which fewer than 10 articles were identified was categorised into ‘Other’. For standardisation purposes, the nomenclature of fetal calf serum (FCS) was aligned with fetal bovine serum (FBS). However, heat-inactivated serum was maintained as a distinct category, as not all serums undergo heat-inactivation during the manufacturing process, as indicated by manufacturer’s documentation (13).

**Figure 3.**
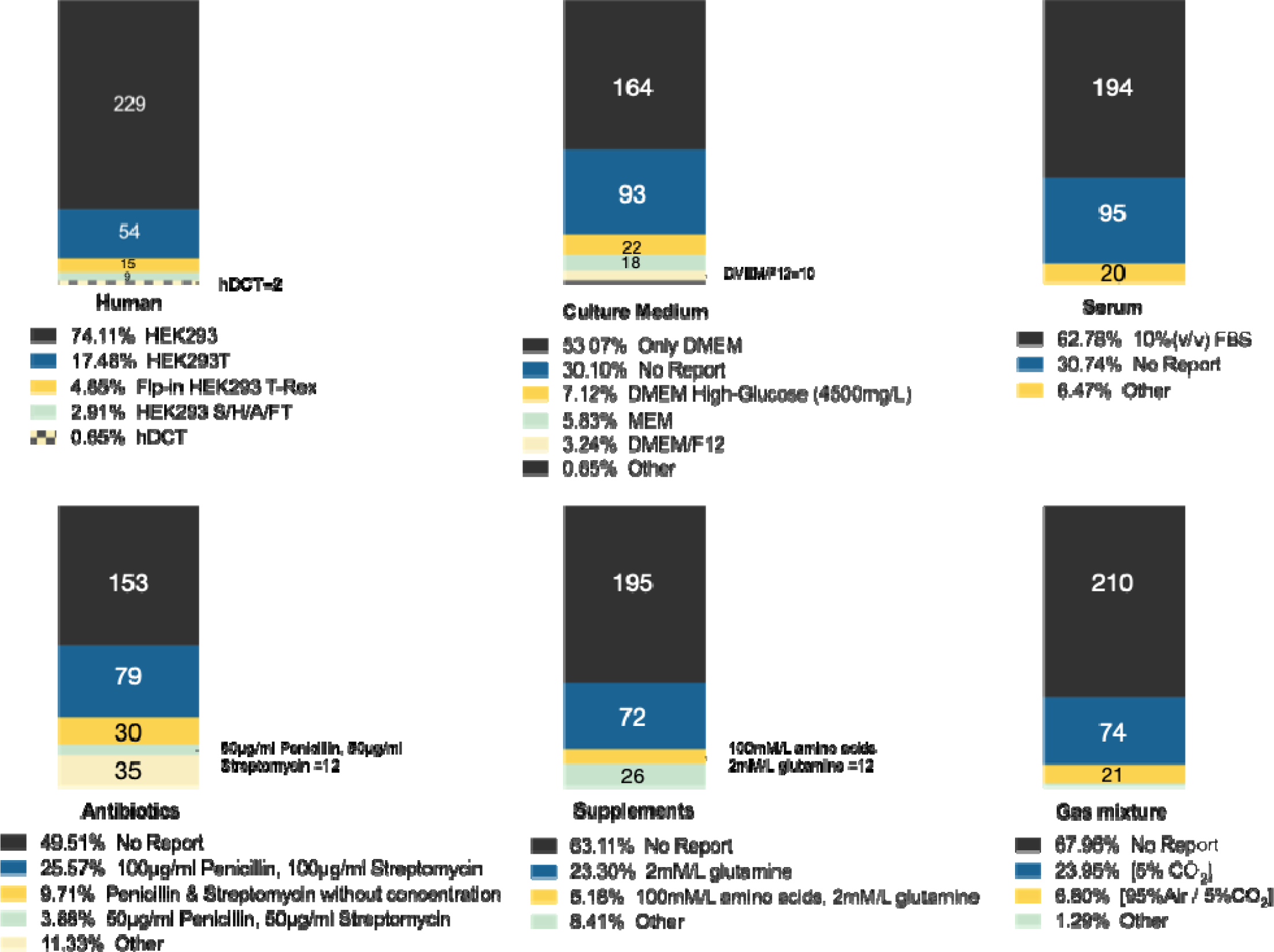
Overview of cell culture condition in HEK293 cell lines and hDCT. The culture condition was categorised into the choices of cell line, culture medium, serum, antibiotics, supplements and gas mixture. The total number of all the charts referred to the total number of culture conditions mentioned in the selected articles, equalling to 309. Any option within each category that used less than 10 times in the selected articles would be sorted into “Other”. **Culture Medium:** MEM= Eagle’s Minimum Essential Medium, DMEM/F12= Dulbecco’s Modified Eagle Medium: Nutrient Mixture F-12. DMEM/F12 was mixed at 1:1(v/v) ratio. **Serum:** All the fetal calf serum (FCS) identified in the articles were renamed to fetal bovine serum (FBS). **Antibiotics:** The concentration of antibiotics was converted to μg/mL.

The majority of reviewed studies used Dulbecco’s Modified Eagle’s Medium (DMEM) and 10% (v/v) FBS for cell culturing. There was significant variability in the selection of antibiotics and other supplements. The combination of 100 μg/mL penicillin and 100μg/mL streptomycin emerged as the predominant choice, cited in 81 articles, followed by a combination of 50 μg/mL penicillin and 50 μg/mL streptomycin, identified in 12 papers. Within the subset of 116 articles that reported on supplement use, 2mM/L glutamine was identified as the most commonly employed supplement in 74 studies. The secondary preference was a mix of 100mM/L amino acids and 2mM/L glutamine. Other supplements that were frequently included are glucose and sodium pyruvate, although their concentrations varied. The incubation temperature for culture was consistently reported at 37°C across all studies. Notably, a significant number of articles did not provide detailed information on culture conditions, with omissions in methodology reporting ranging from 30% for culture medium and serum specifics to 68% for gas mixture details, highlighting a gap in comprehensive documentation.

In response to the considerable number of studies that did not fully disclose cell culture conditions, **Table 1** presents the three most prevalent cell culture recipes for each cell line, based exclusively on articles that comprehensively reported on the culture medium, serum, antibiotics, supplements, and their concentrations. This ranking was determined by the frequency of each combination’s usage. Across various cell lines, the most common incubation environment was a humidified atmosphere of 95% air and 5% CO_2_ at 37℃.

**Table 1.** The 3 most common cell culture conditions used in different HEK293 cell lines and in hDCT. Only articles that fully reported culture medium, serum, antibiotics and supplements with concentration stated were included into tabulation. The preference was ranked by the frequency of use. The number in bracket represented (number of papers used this recipe / number of articles that fully reported the culture condition). The preferred incubation environment was 95% air /5% CO_2_ at 37℃ in all cell lines.

| Cell line | Recipe 1 | Recipe 2 | Recipe 3 |
| --- | --- | --- | --- |
| HEK293 | DMEM, 10% FBS, 100µg/mL penicillin, 100µg/mL streptomycin, 2mM/L glutamine (66/116) | DMEM/F12, 10% FBS, 50µg/ml penicillin, 50µg/ml streptomycin, no supplement reported (9/116) | MEM, 10% FBS, 50µg/ml penicillin, 50µg/ml streptomycin, no supplement reported (6/116) |
| HEK293T | DMEM, 10% FBS, 100µg/mL penicillin, 100µg/mL streptomycin, 2mM/L glutamine (21/24) | MEM, 10% FBS, 2mM/L penicillin, streptomycin, no supplement reported (3/24) |  |
| Flip-in HEK293T | DMEM high glucose (4500mg/L), 10% FBS, 200µg/ml hygromycin, 15µg/ml blasticidin, 2mM/L glutamine (8/16) | DMEM, 10%FBS, 5µg/ml blasticidin, penicillin or streptomycin or zeocin or hygromycin, no supplement reported (7/16) | DMEM, 10% FBS, 100µg/mL penicillin, 100µg/mL streptomycin, no supplement reported (1/16) |
| Concentration of tetracycline or doxycycline used for transfection induction | 1 µg/mL tetracycline or doxycycline for induction. (11/14) | 2 µg/ml tetracycline for induction (1/14) | 0.1 µg/ml tetracycline for induction (1/14) |
| HEK293A/S/H/FT | DMEM, 10% FBS, 100µg/mL penicillin, 100µg/mL streptomycin, 2mM/L glutamine (5/9) |  |  |
| hDCT | DMEM/F12, 10% FBS, 100µg/mL penicillin, 100µg/mL streptomycin, 10ng/mL in human recombinant epidermal growth factor, 36 ng/mL dexamethasone, 2 ng/mL triiodothyronine, insulin-transferrin-selenium at 1:100 into medium (1/2) | DMEM/F12, 10% FBS, 100µg/mL penicillin, 100µg/mL streptomycin, insulin-transferrin-selenium at 1:100 into medium, 2.5 µg/mL Nicotinamide at 1:5000 into medium, 500µg/mL hydrocortisone at 1:2500 into medium (1/2) |  |

Specifically, for the HEK293, HEK293T, and other related HEK293 variants, the predominant cell culture formula consisted of DMEM, supplemented with 10% FBS, 100 μg/mL penicillin, 100 μg/mL streptomycin, and 2 mM/L glutamine. For experiments involving Flp-In HEK293 T-REx cells, the preferred medium was DMEM with high glucose (4500 mg/L), 10% FBS, 200 μg/mL hygromycin, 15 μg/mL blasticidin, and 2mM/L glutamine, supplemented with 1 μg/mL tetracycline or doxycycline to induce T-Rex driven gene expression.

The culture supplement for human distal convoluted tubule (hDCT) cell cultures diverged from other lines, incorporating 10 ng/mL human recombinant epidermal growth factor, 36 ng/mL dexamethasone, 2 ng/mL triiodothyronine, and insulin-transferrin-selenium in a 1:100 ratio to the medium. Alternatively, nicotinamide and hydrocortisone were added into the culture. This unique selection of supplements for hDCT cultures likely aims to mimic physiological endocrine and paracrine signalling, thereby providing an optimised context for studying the nuanced functions and responses of these renal tubular cells under conditions that closely replicate their *in vivo* environment.

### Culture Condition: Murine Cell Lines

Culture conditions were sorted into culture medium, serum, antibiotics, supplements and gas mixture (**Figure 4**). The culture conditions for three mDCT cell lines demonstrate a high level of uniformity, particularly in the choice of culture medium, with Dulbecco’s Modified Eagle Medium/F12 (DMEM/F12) being the preferred medium in the vast majority of cases. This consistency is not reflected in the reporting of serum, antibiotics, and supplements, where there is a notable lack of comprehensive documentation.

**Figure 4.**
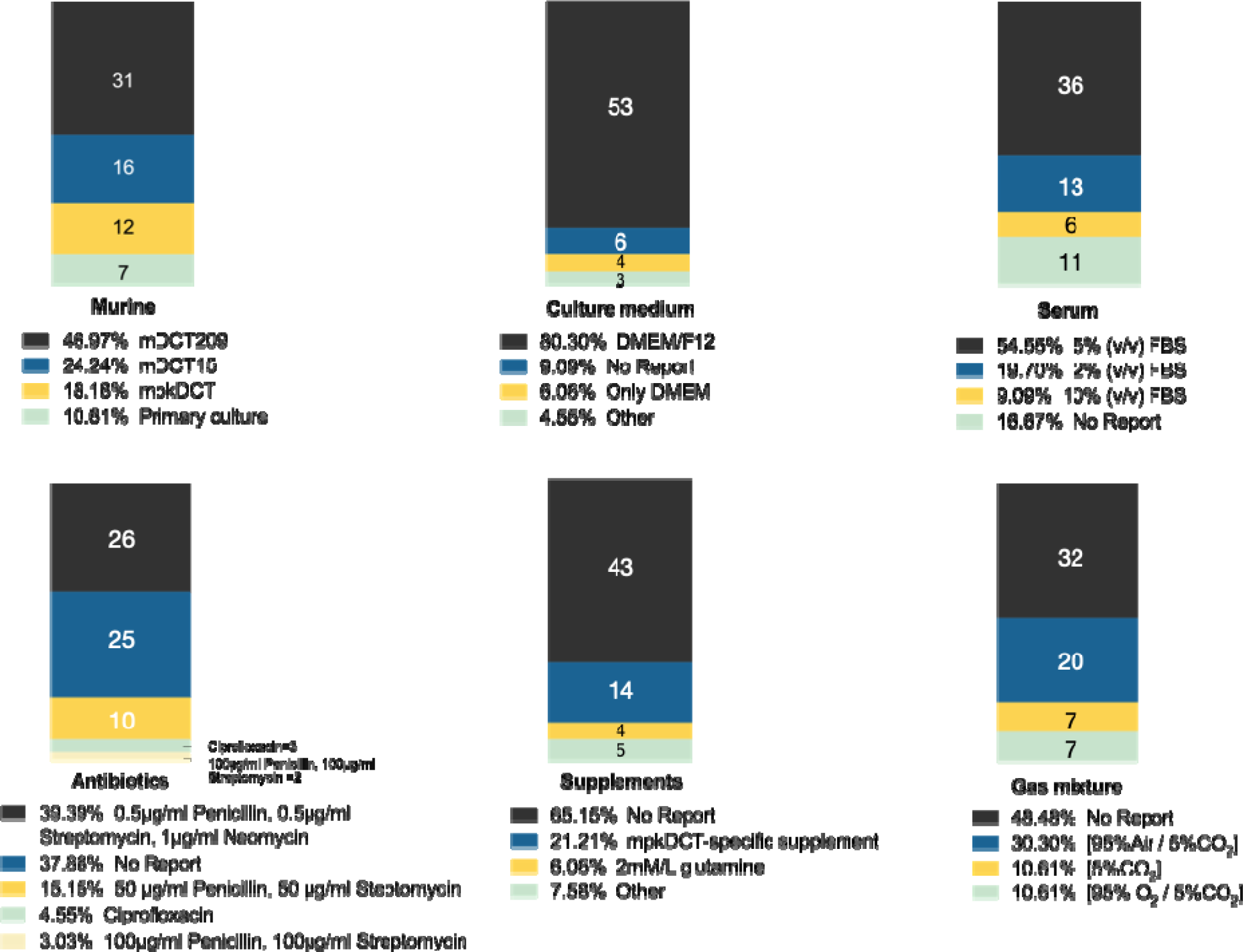
Overview of cell culture condition in murine DCT cell lines. Total conditions included in analysis n=66. **Culture medium:** Other medium that used in culture include Eagle’s Minimum Essential Medium (MEM), DMEM High-Glucose (4500mg/L) and DMEM/F12. **Antibiotics**: the concentration and unit of antibiotic mixture was standardised to μg/mL. **Supplements:** mpkDCT had two sets of culture conditions exclusively used in mpkDCT studies and full detail of recipes can be found in **Table2**.

For the serum supplementation in mDCT cell line cultures, approximately half of the documented studies reported using a 5% FBS, regardless of the serum choice. The second preferred concentration was 2% serum, which accounted for about 20% of the cases. The remaining 10% of studies opted for a higher concentration, utilising 10% serum for their mDCT cell cultures. This distribution suggests that while there is a predominant trend towards lower serum concentrations, there is still considerable variability in serum use among the studies. Among those that specified antibiotics, a combination of 0.5 μg/mL penicillin, 0.5 μg/mL streptomycin and 1μg/m neomycin was most common, appearing in 38.18% of studies, and a higher concentration combination of 50 μg/mL each was used less frequently. Regarding supplements, a significant 61.15% of papers did not report on this aspect. For those that did, 2 mM/L glutamine was a common addition, followed by a mix of 5mM glucose & 5mM glutamine, and the use of HEPES buffer or 1g/L glucose & 1mM sodium pyruvate as other supplement options.

**Table 2** summarised the prevalent culture conditions for mDCT cells, with an inclusion criterion focused on articles that thoroughly documented the culture medium, serum, antibiotics, and supplements. The culture protocol for the mpkDCT cell line exhibited remarkable uniformity, following the standardised combination established by the cell line’s inventor, Professor Alain Vandewalle. Nonetheless, this review underscores substantial deficiencies in the methodological reporting across murine cell line studies, especially in the areas of serum and supplement use. The frequency of such reporting gaps poses significant challenges to the scientific community, as it impairs the ability to replicate and rigorously compare experimental outcomes across the discipline.

**Table 2.**
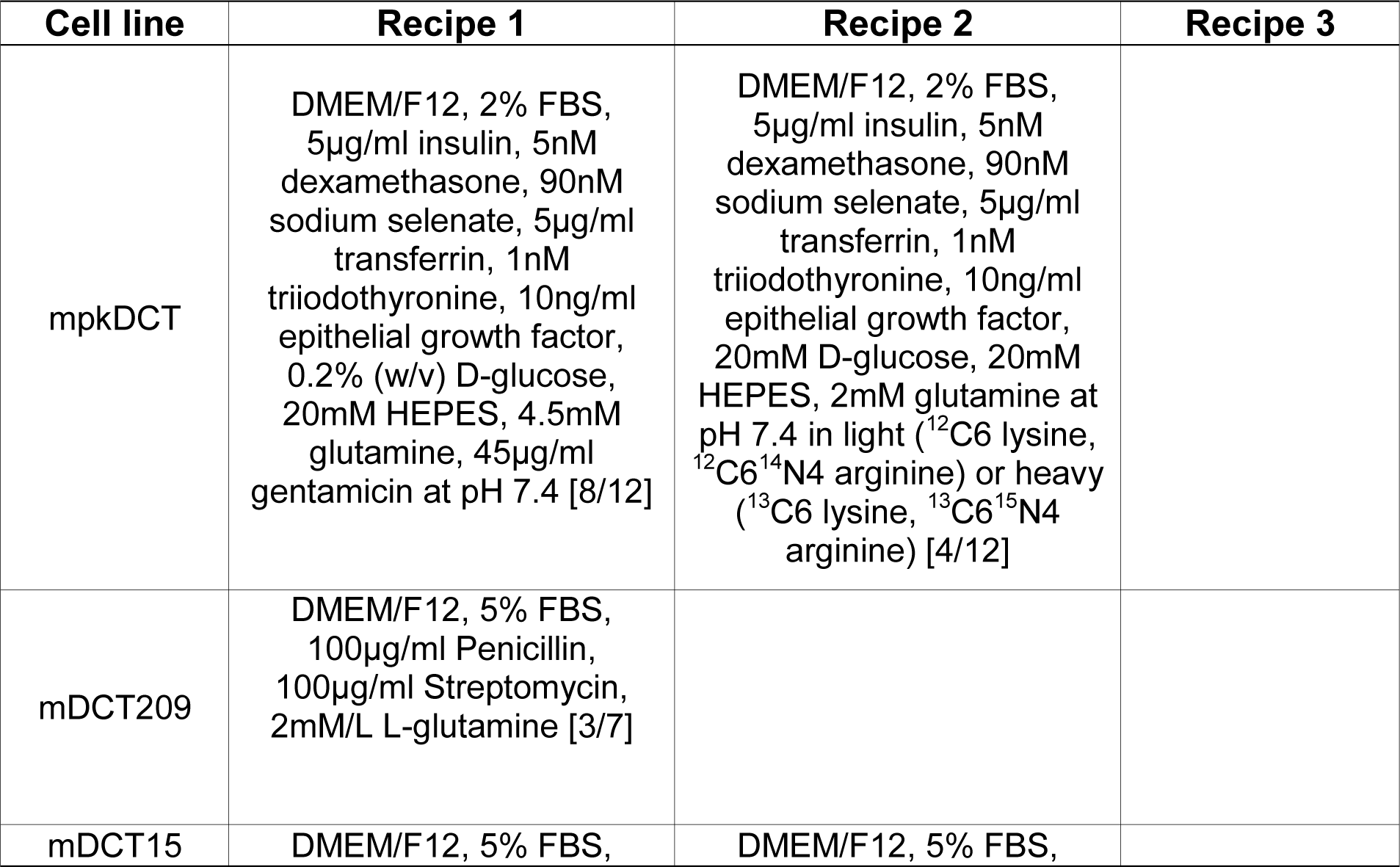
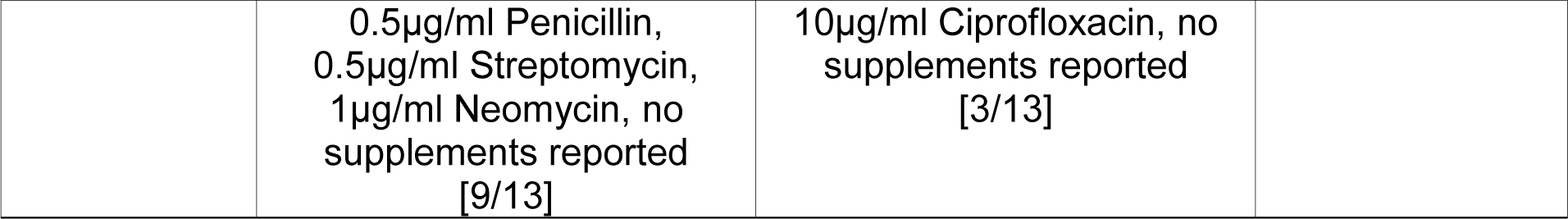
The 3 most common cell culture conditions used in different mDCT cell lines. Only articles that fully reported culture medium, serum, antibiotics with concentration stated were included into tabulation. The preference was ranked by the frequency of use. The number in bracket represented [number of papers used this recipe / number of articles that fully reported the culture condition]. The preferred incubation environment was 95% air /5% CO_2_ at 37℃ in all cell lines.

### Culture Condition: Canine Cell Lines

There is a notable limitation in the breadth of available literature for culturing MDCK and its derivatives. Only a select number of studies met the criteria for inclusion, and even fewer provided a full account of their culture conditions. Given this scarcity of comprehensive data, we have endeavored to summarise all pertinent culture information within **Table 3** (14–26).

**Table 3.**
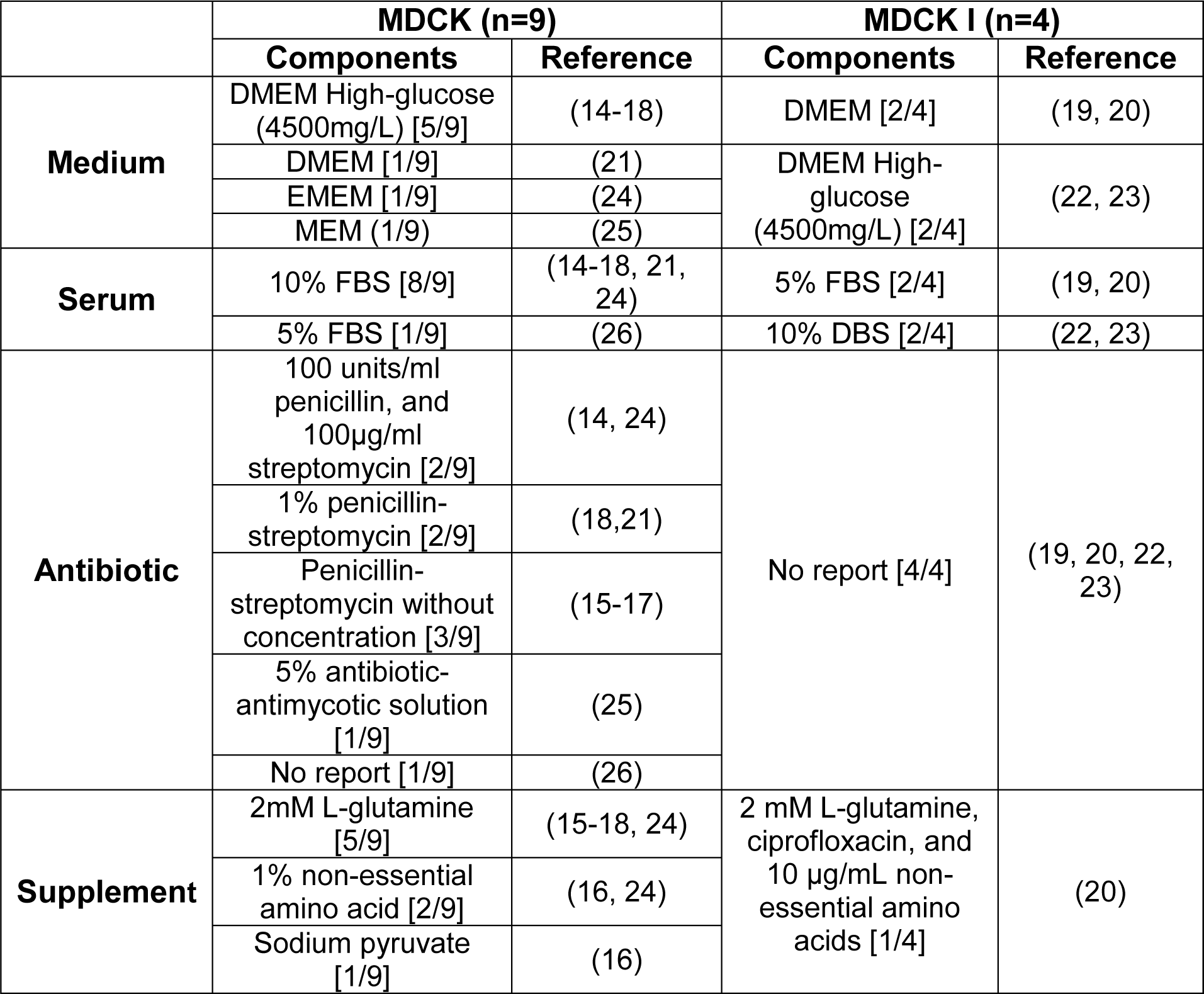
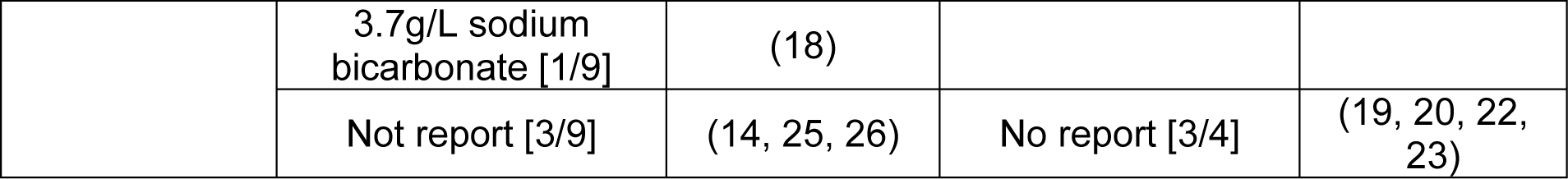
Overview of cell culture conditions in MDCK and MDCK I cell lines. The culture conditions were categorised into the choices of cell line, culture medium, serum, antibiotics and supplements. Only articles that fully documented the choice of culture medium and serum were included into table. The number in bracket represented [number of papers used this component / number of articles that fully reported the culture condition]. The preferred incubation environment was 95% air /5% CO_2_ at 37℃ in all cell lines. **EMEM=** Eagle’s Minimum Essential Medium.

### Transfection Methods in HEK293 Lines

The versatility of HEK cells and their derivatives is underscored by the high volume of transfection activities reported. This suggests an inherent adaptability or ease of transfection associated with these cell lines, making them a popular choice for genetic manipulation studies. To manage the extensive data on transfection activities, an arbitrary threshold was established for inclusion in the analysis: only transfection methods reported at least 50 times were assigned a distinct identifier; those with fewer instances were collectively categorised under ‘Others.’ This approach facilitated a more streamlined and focused analysis. The three transfection reagents that emerged as the most frequently utilised were Lipofectamine 2000, FuGENE 6, and polyethyleneimine, with 281, 96, and 60 documented uses, respectively, as detailed in **Figure 5**. Moreover, it was observed that detailed reporting on transfection efficiency was remarkably infrequent, with only 6 out of 301 articles addressing this metric. Furthermore, the documentation of cell confluency at the time of transfection and the nature of the transfection, whether transient or stable, was inadequately reported, with only 10% of studies mentioning confluency and 23% specifying the transfection state. This lack of comprehensive reporting on critical aspects of the transfection process hampers the ability to fully assess the methodologies and their efficacy.

**Figure 5.**
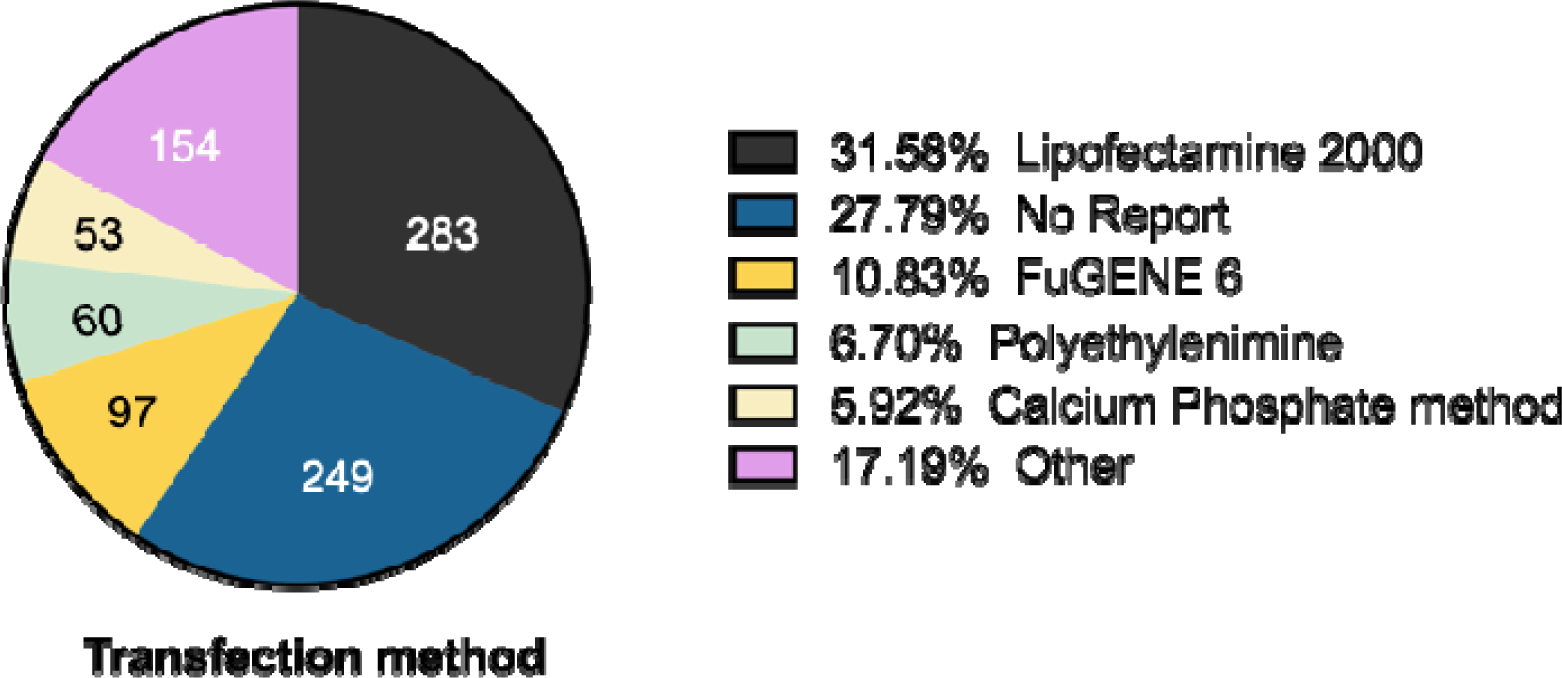
Overview of transfection reagents used in HEK293 and derivatives. The total number of transfection activities=898. The threshold of display was set at 50 times for transfection reagent and any reagent used less than the threshold would be grouped into “Other”. Percentage was calculated as times of transfection activity divided by total number of transfection reagent used.

### Transfected Genes in HEK293 Lines

Figure 6 presents a heat map illustrating the frequency of wild type and mutant gene transfections into HEK293 cells and their derivatives. A total of 181 wild type and 57 mutant genes have been introduced into these cell lines. For clarity in visualisation, only wild type genes with a minimum of three transfections were included in the heat map, showcasing 51 genes. All mutant genes were displayed without a threshold. The most frequently transfected wild type genes were the Potassium Inwardly Rectifying Channel Subfamily J member 1 (*KCNJ1*, also known as Kir1.1 or ROMK), WNK1, and WNK4. For mutant genes, WNK4 and the *SLC12A3*, were the most commonly used. Despite the widespread use of HEK293 cells and their derivatives as a model for studying DCT physiology, it is important to note that they do not inherently express many of the key channels and transporters characteristic of native DCT cells. This necessitates the frequent transfection of such genes to render them suitable for DCT-specific research, as reflected in the transfection data presented in the figure.

**Figure 6.**
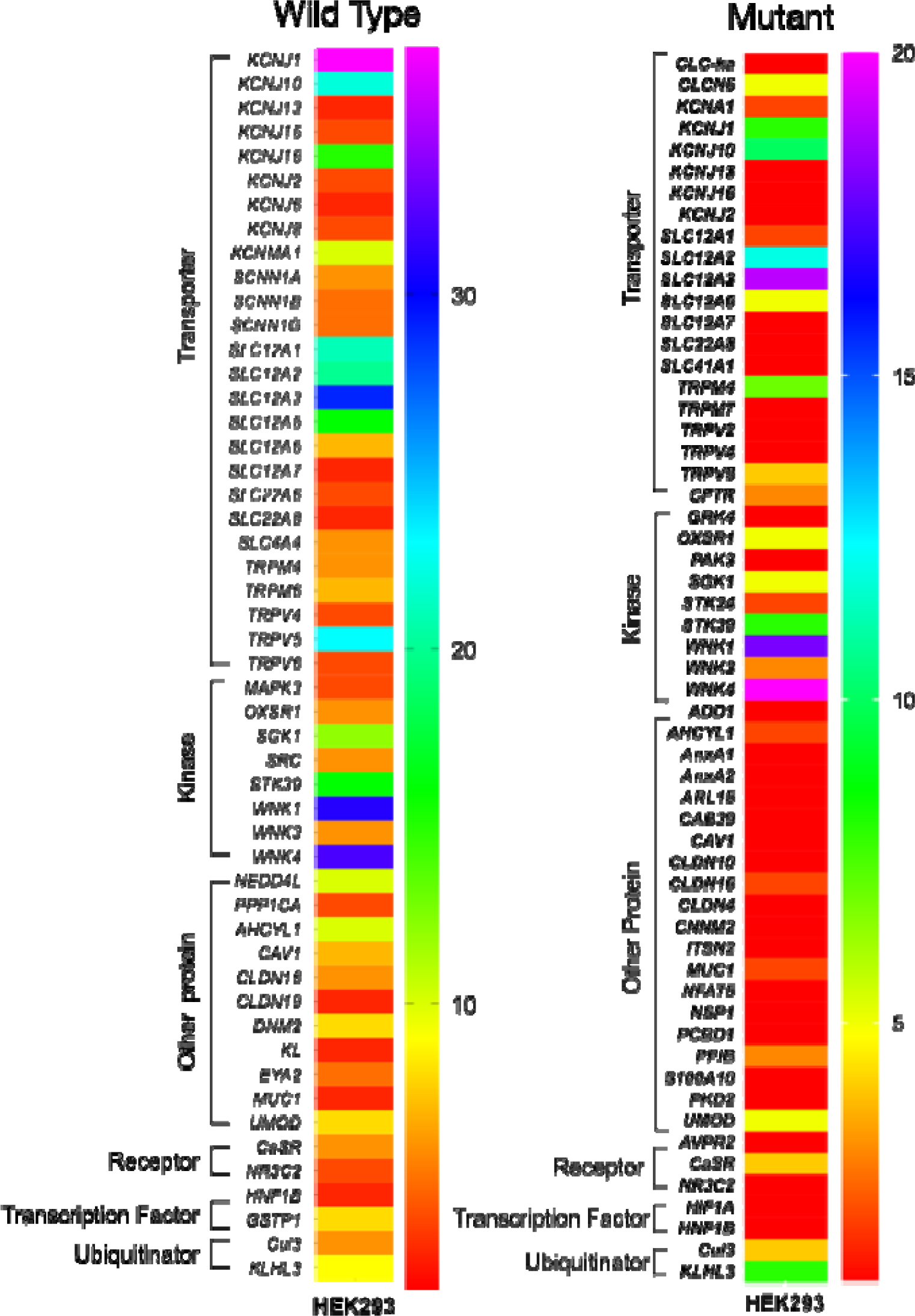
Heat map of the frequency of wild type and mutant gene transfections into HEK293 and derivatives. Color-coded values represent the total frequency of transfection in to any HEK293 cell lines of this gene. Genes were clustered by function. **Left:** Heat map of wild type genes transfected into HEK293 and derived cell lines. Threshold of display was set at transfected for more than 3 times, giving 51 wild type gene shown in this heat map. **Right**: Heat map of mutant genes transfected into HEK293 and derived cell lines. No threshold was set for display, giving 57 mutant genes transfected shown.

### Endogenous Genes in DCT Cell Lines

A thorough understanding of the endogenous gene expression profiles of cell lines is important, as it sheds light on the inherent cellular functions and potential constraints of these lines. Figure 7 offers a heat map that delineates the types and frequencies of endogenous gene expressions documented in the literature for HEK293 cells and their derivatives. This heat map includes 34 genes that have been validated through more than one study or method. Prominently, two housekeeping genes, α/β-actin and GAPDH, stand out as the most commonly detected, with 32 and 19 instances, respectively. The heat map highlights the prevalence of genes such as WNK4, STK39, and OXSR1, signifying the HEK293 line’s widespread adoption as a model for probing the WNK-SPAK/OXSR1 pathway. This is in spite of the HEK293’s lack of many critical downstream transporters involved in this pathway; a potential area for model enhancement. In contrast, mDCT models, as more recent entrants into the field, offer a targeted examination of the DCT region. As shown in Figure 8, the mDCT cell lines demonstrate a consistent endogenous expression of the *SLC12A3*, evidenced by its prevalent detection in mDCT209, mDCT15, and mpkDCT, with expression counts of 29, 23, and 27, respectively. The repeated identification of *SLC12A3* attests to the strength of these models for DCT-specific research. Moreover, the presence of other vital transporters, such as TRPV5, TRPM6, and members of the PMCA family, concurs with the DCT’s distinct ion transport functions. Notably, the ENaC channel’s presence in mDCT15 and mpkDCT lines further corroborates their relevance for studies into DCT-related sodium reabsorption. The regular reporting of WNK4, STK39, and OXSR1 in these lines reinforces their value in elucidating the complexities of the WNK-SPAK/OSR1 signaling pathway, a key player in ion homeostasis and blood pressure regulation.

**Figure 7.**
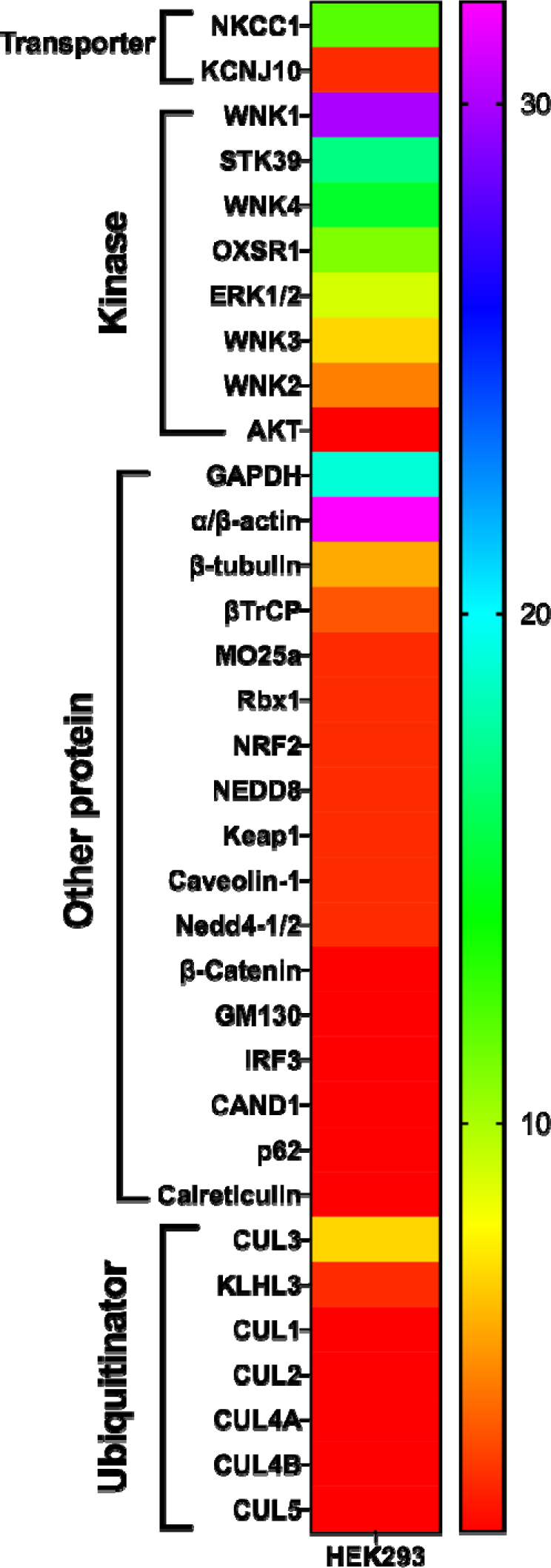
Heat map of the times of endogenous gene detected in HEK293 and derivatives. Color-coded values represent the total time of this gene identified in any HEK293 cell lines. Genes were clustered by function. Threshold of display was set at detected for more than 2 times or by more than 2 methods. There are 34 genes presented in total, of which α/β-actin, WNK1 and GAPDH are the top three detected expression with32, 30 and 19 times reported respectively.

**Figure 8.**
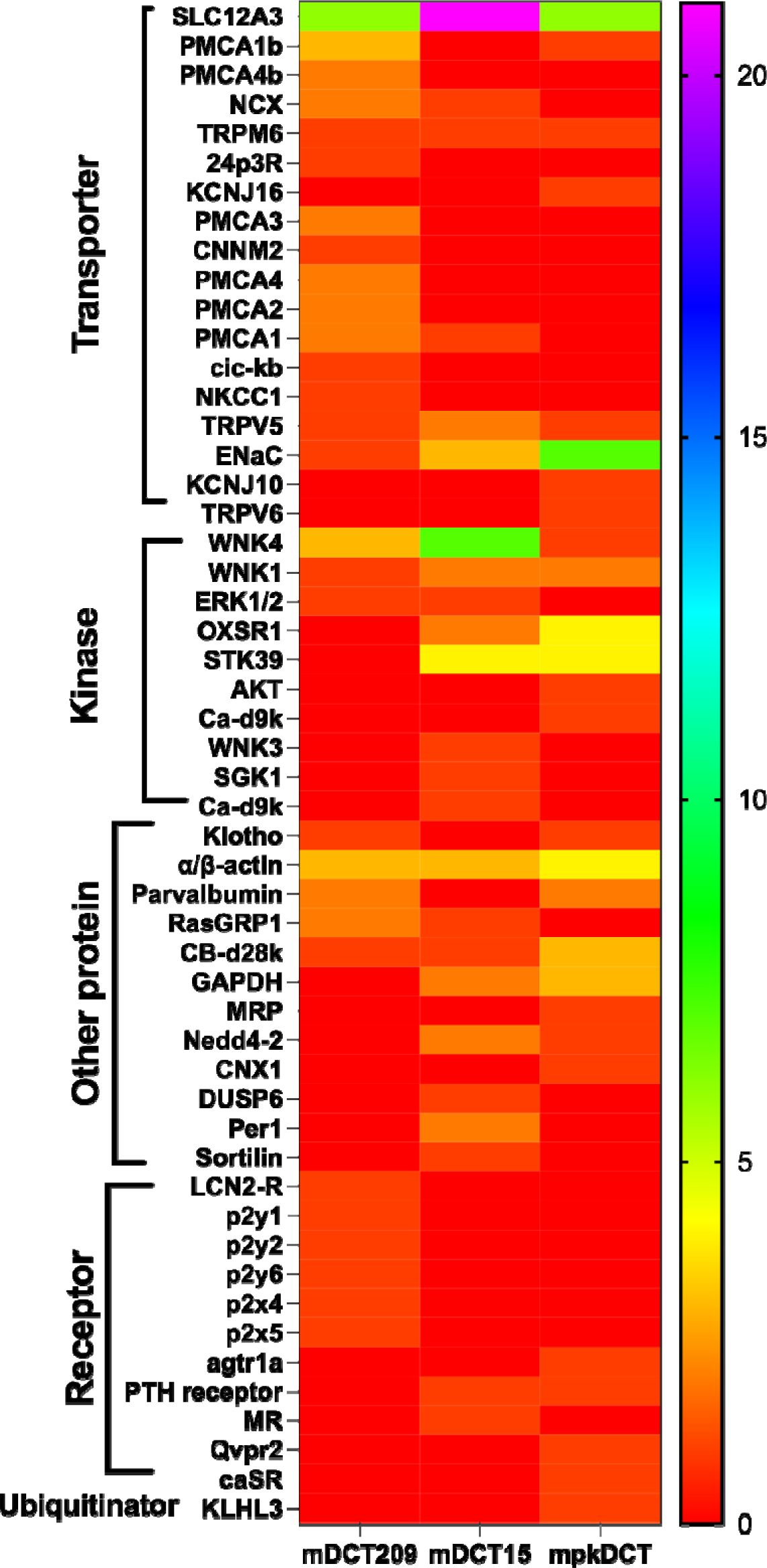
Heat map of the times of endogenous gene detected in mDCT209, mDCT15 and mpkDCT. Color-coded values represent the total time of this gene identified in the cell line. Genes were clustered by function. There are 29, 23 and 27 genes natively expressed in mDCT209, mDCT15 and mpkDCT, respectively.

## DISCUSSION

While the influence of the Renin-Angiotensin-Aldosterone System (RAAS) on blood pressure regulation through renal mechanisms has been recognised for many years, it is only with recent advances in understanding the WNK-SPAK signalling pathway and the pathology of Familial Hyperkalemic Hypertension that the critical role of the DCT in maintaining blood pressure homeostasis has been underscored. Given the limitations in conducting in vivo studies, the development and use of appropriate *in vitro* cellular models have become indispensable tools in the exploration of these mechanisms.

### Lineage of DCT Cell Lines

Figure 8 delineates the overview of existing DCT models discussed in this review. The original HEK293 cell line was immortalised from the embryonic kidney of a female fetus by Alex Van der Eb in 1977 at the University of Leiden, Netherlands (27). This initial cell line has since given rise to over 12 derivative cell lines (28). The proliferation of these derived cell lines into commercial products and their extensive use in both laboratory and industrial settings have introduced complexities, particularly in nomenclature and lineage documentation (29). The diversity in the naming of HEK293 strains and the often ambiguous documentation of their derivation pose challenges in accurately tracing their origins. Genomic and transcriptomic analyses have revealed significant variations in the chromosome numbers among HEK293 cells obtained from different commercial sources, indicating karyotypic drift from the original cell line. This drift likely results from repetitive and non-standardised cultivation practices across various settings (30). Such genomic instability has led to noticeable changes in the genotype and phenotype of the cell line and its derivatives, manifesting during processes such as clonal isolation, mutation, and expansion (10, 30, 31). Moreover, the HEK293 line was derived from an entire embryonic kidney, without specific isolation from the distal convoluted tubule (DCT) or any other kidney segment. The embryonic origin of these cells suggests that HEK293 cells may not accurately reflect the physiology of adult kidney cells. Intriguingly, recent studies have shown that the transcriptome of HEK293 cells more closely resembles that of neuronal cells, proposing an adrenal or neuroendocrine origin rather than a renal one (31). HEK293 cells and their derivatives are highly favored in transfection studies, as evidenced by the analysis of 301 research papers, which identified a total of 898 transfection events. These involved the transfection of 181 wild type and 57 mutant genes at least once into HEK293 or its derivatives, predominantly through transient methods. Key wild type genes such as *KCNJ1* (encoding Kir1.1/ROMK/ROMK1), WNK1, WNK4, and *SLC12A3* emerged as the most commonly transfected, with each being introduced approximately 30 times. Mutant versions of *WNK4, SLC12A3*, and *WNK1* were also frequently used in these cell lines. The high volume of transfection activities suggests the HEK293 cell lines’ favorable characteristics for transfection, highlighting their role as efficient vectors for gene expression studies. However, this extensive reliance on transfection might also suggest that HEK293 cells and their derivatives fall short as direct models for DCT research, necessitating the introduction of exogenous genes to compensate for the absence of native DCT gene expression. Presently, there is a noticeable gap in detailed profiling of the HEK293 cell line’s endogenous gene expression. This review identified the endogenous presence of several genes in HEK293 or its derivatives, including Potential Cation Channel Subfamily M Member 4 (*TRMP4*), Calcium Binding Protein 39 (*CAB39*), Intersectin 1 (*ITSN1*), Serine/Threonine Kinase 39 (*STK39*), Caveolin-1, *OxSR1*, *WNK1* and *WNK4* (16, 32–39). Contrastingly, crucial genes associated with solute transporters, such as *SLC12A3, TRPM6, KCNA1* (encoding for the Kv1.1 potassium channel), *SCNN1A* (encoding the ENaC alpha subunit), *KCNJ10* (encoding Kir4.1), and *KCNJ16* (encoding Kir5.1), are not natively expressed in HEK293 cells, necessitating their transfection for study. For example, the DCT plays an exclusive role in transcellular magnesium reabsorption with *TRPM6*, a process tightly regulated and essential for maintaining magnesium homeostasis (40), while HEK293 does not natively expressed. The DCT is characterised by its natively low chloride concentration, a feature that significantly influences the electrochemical gradient and drives the reabsorption of sodium and calcium ions. This low chloride environment is essential for the proper functioning of various transporters and channels within the DCT (41, 42). Significant efforts have been made to modify HEK293 cell lines to replicate the potassium sensing and transport characteristics intrinsic to *in vivo* DCT cells, addressing the inherent limitations of HEK293 cells in this regard. HEK293 lines lack the native capability to mimic the DCT’s specific responses to changes in extracellular potassium concentrations. While this process presents considerable challenges, the advancements signal a move towards more representative models for studying renal potassium regulation and DCT cell function (43). This discrepancy underscores the need for transfection to introduce key DCT functionalities into HEK293 cell-based models, pointing to their limitations in mimicking native DCT gene expression profiles (44).

**Figure 8.**
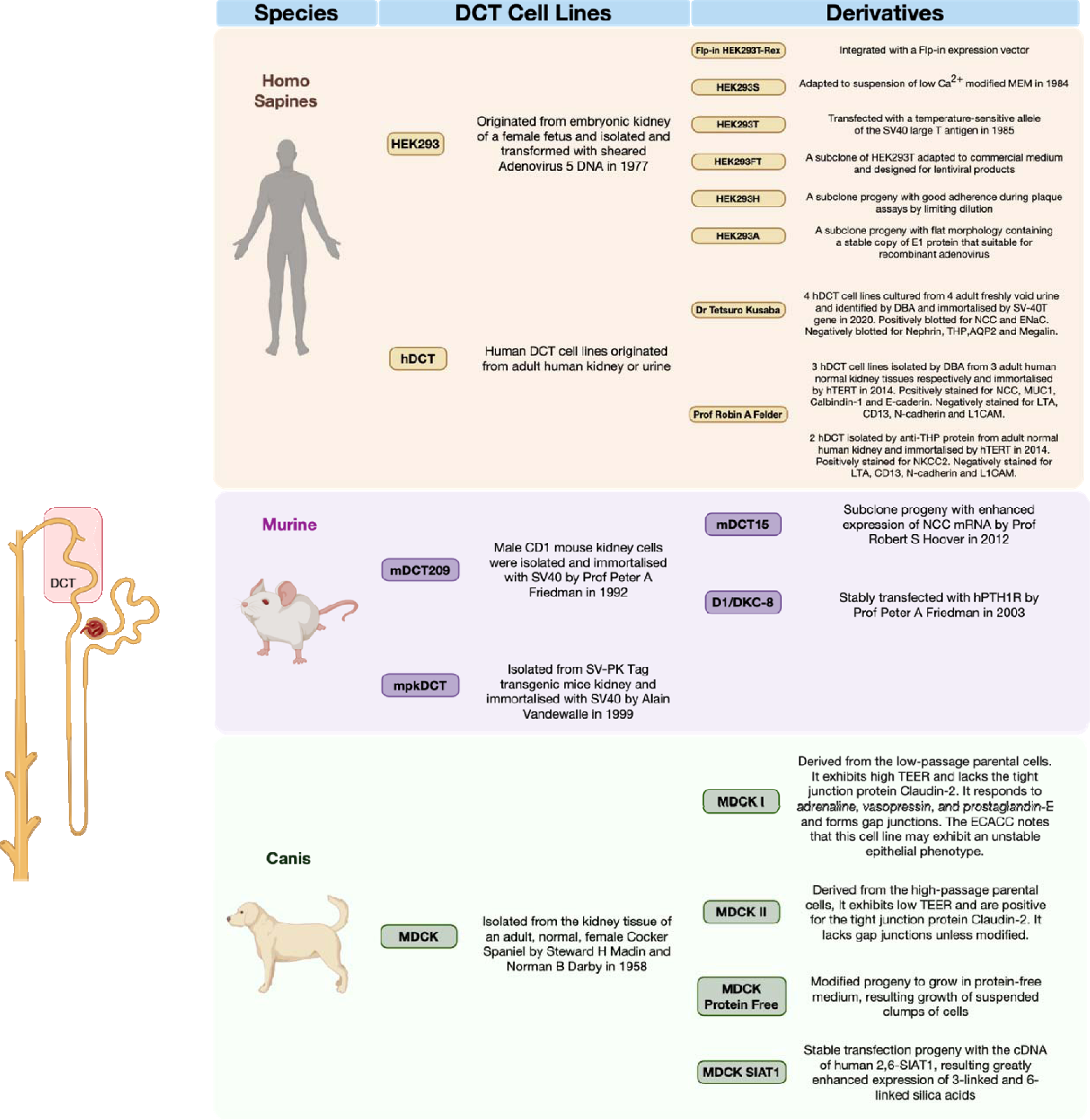
An overview of the development and linage relationships of human renal models that discussed in this study. DBA= dolichos biflorus agglutinin. hTERT= human terminal transferase. TEER= Trans-epithelial/trans-endothelial electrical resistance. hPTH1R= human parathyroid hormone receptor. The years stated in the figure refer to the year of cell line publicised on journals, not necessary the year of invention.

mDCT cell lines are highly regarded for their clearly defined origins and the preservation of essential ion transport processes and regulatory mechanisms. Among them, mDCT209, mDCT15, and mpkDCT are the most widely utilised models, while primary cultures also being a preferred method for investigating DCT functionality, albeit to a lesser extent. The mDCT209 cell line, established in 1992 by Professor Peter A. Friedman through the isolation of DCT cells from CD1 mice, marked the inception of the first immortal murine DCT cell model. It is known for naturally expressing NCC and the parathyroid hormone receptor (PTHR) (45). Following this, the mpkDCT cell line was derived in 1999 by Professor Alain Vandewalle from the kidney of an adult transgenic mouse, showcasing the expression of calcium channels like TRPV5 and TRPV6, and has been instrumental in studies of calcium transport (46, 47). Addressing the absence of a mammalian DCT cell model with consistently high expression of the NCC, Professor Robert S. Hoover and his team developed the mDCT15 cell line. This new line was derived from the existing mDCT209 line through a process of screening and subcloning cells that exhibited the highest levels of NCC mRNA, as determined by real-time PCR (48). The variation in methodologies employed for isolating the initial primary material has resulted in each of the three mDCT cell lines—mDCT209, mDCT15, and mpkDCT—exhibiting unique gene expression profile and representing distinct regions of the DCT. This diversity is crucial for researchers when selecting the most suitable cell line for their specific experimental needs, underscoring the importance of considering these differences in genetic characteristics and regional representation in experimental design. The mDCT209 cell line has been validated to express *SLC12A3* (49, 50) and parvalbumin (51), indicative of a heterogeneous population that encompasses both early (DCT1) and late (DCT2) segments. This cell line has facilitated seminal research on the WNK signaling pathway (50, 52) and the hormonal regulation by aldosterone (53, 54). Additionally, it has been instrumental in studying the impact of phosphoric acid depletion on magnesium ion reabsorption (55), a process potentially mediated by the TRPM6 channel expressed in these cells (56). As the most established and extensively utilised cell line, mDCT209 has undergone numerous passages across various laboratories, which may have contributed to genetic and phenotypic variations (57), including a notable heterogeneity in NCC expression at the single-cell level (48). This variability underscores the absence of karyotype stability reports for this, as well as other DCT cell lines. To address the limitations of mDCT209, Ko et al. developed the monoclonal mDCT15 cell line in 2012, directly derived from mDCT209 (48). This cell line embodies the characteristics of DCT2 cells more accurately, with a robust expression of functional NCC, WNK, SPAK, OXSR1, NEDD4-2, ROMK and SGK1, alongside another DCT2 marker, the epithelial sodium channel (ENaC), with all its three subunits confirmed (58–63). Meanwhile, the mpkDCT cell line was initially established to explore renal calcium transport mechanisms, expressing additional key calcium transporters and binding proteins such as TRPV5, TRPV6, Calbindin-D9k, and Calbindin-D28k, that are exclusively found in DCT2 (46, 46, 64–66). Distinguishing between DCT1 and DCT2 in research models is crucial due to their distinct roles in renal electrolyte handling, with DCT1 focusing on sodium and chloride reabsorption and DCT2 on calcium, magnesium, and potassium balance (67). This differentiation allows for precise studies relevant to conditions like hypertension and electrolyte imbalances. Moreover, segment-specific responses to hormones and medications underscore the importance of accurate modeling for translational insights into kidney function and related disorders. For the available murine models, as shown in **Figure 9,** the mDCT209 cell line serves as a versatile model, displaying markers of both DCT1 and DCT2 cells, and stands as the most adaptable option currently available for DCT research. In contrast, both mpkDCT and mDCT15 exhibit phenotypes more characteristics of DCT2. This delineation highlights a clear gap in the availability of an immortalised DCT1-like cell line that could more distinctly differentiate between DCT1 and DCT2 in terms of function and regulation, suggesting an ongoing need for more specialised models.

**Figure 9.**
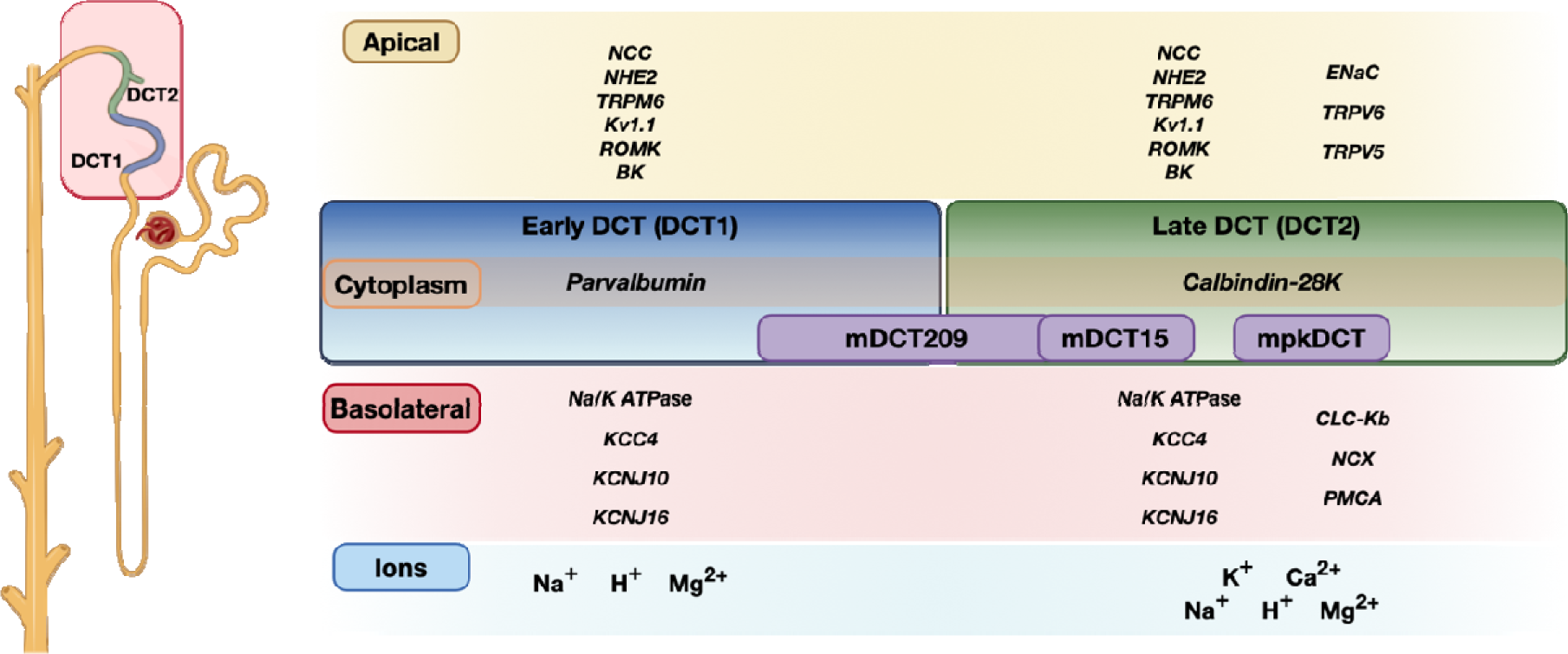
The schematic illustrates the distribution of DCT1 and DCT2 markers and how they correlate with the phenotypes of various mDCT cell lines. Transporters and markers specific to human DCT1 and DCT2 are categorised on the left and right sides of the schematic, respectively. The placement of mDCT cell lines within this framework serves as a conceptual representation to show the extent to which each cell line mirrors the native phenotypes of DCT1 and DCT2. Through comparative analysis of endogenous gene expression between the mDCT cell lines and native mammalian DCT segments, it has been determined that mDCT15 and mpkDCT exhibit phenotypes that align closely with DCT2. Conversely, mDCT209 presents a composite phenotype that integrates characteristics of both DCT1 and DCT2, albeit with a slight inclination towards a DCT1-like phenotype. This schematic aids in visualising the relative positioning of each cell line within the spectrum of DCT functionality, offering insights into their suitability as models for specific segments of the distal convoluted tubule

Species differences pose a significant challenge in utilising DCT models and animal models for research that aims to extrapolate findings to humans. The molecular cloning of the human NCC has revealed the existence of three distinct isoforms resulting from alternative splicing. Of these, isoforms NCC_1_ and NCC_2_, which consist of 1030 and 1029 amino acids each, are exclusively identified in humans and higher primates and functionally distinct, whereas NCC_3_ with 1021 amino acids (68). The splicing variants of the NCC exhibit discrepancies in naming between the NCBI and UniProt databases. The **Table 4** summarises these differences to clarify the nomenclature used across these platforms. This specificity highlights a potential limitation in the translational applicability of findings from mDCT models to human physiology. Studies on the serine/threonine kinase SPAK have identified differences in SPAK isoform expression between human and mouse kidney tissues. Although human tissues express two kinase-deficient proteins, SPAK2 and KS-SPAK, with transcripts similar to those found in murine renal tissue, the relative abundance of these isoforms— distinguished by their molecular weights—differs significantly. These isoforms are also transcribed from a human-specific promoter, indicating that STK39 is subject to species-specific transcriptional regulation and suggesting a potentially unique role in blood pressure regulation across species (69).

**Table 4:** Comparison of NCC Splicing Variant Nomenclature between NCBI and UniProt Databases. The table delineates the divergent naming conventions applied to the splicing variants of the NCC as found in the NCBI and UniProt databases.

| NCBI Name | Amino acids (n) | NCBI Accession | UniProt Name | UniProt Code |
| --- | --- | --- | --- | --- |
| NCC<sub>1</sub> | 1030 | NP_000330.3 | NCC2 | P55017-2 |
| NCC<sub>2</sub> | 1029 | NP_001119579.2 | NCC3 | P55017-3 |
| NCC<sub>3</sub> | 1021 | NP_001119580.2 | NCC1 | P55017-1 |

### Development of human-derived DCT Cell Lines

Given the aforementioned challenges associated with HEK293 family and mDCT models in faithfully replicating human DCT environment, there has been a pronounced emphasis on the development of more accurate, human-derived DCT models for research. This shift towards leveraging human-specific models aims to surmount the limitations observed, enabling a closer approximation of human renal physiology. In 2014, Gildea and colleagues were, to our knowledge, the first to report five novel human DCT cell lines, generate from normal nephrectomy tissues from human adult donors (11). Three hDCT cell lines were isolated using biotinylated *Dolichos Biflorus Agglutinin* (DBA) followed by anti-biotin magnetic nanoparticle-conjugated secondary antibodies. These lines expressed key distal tubule markers, including the NCC, epithelial membrane antigen (MUC1), Calbindin 1, and E-cadherin. In an alternative approach, two hDCT cell lines were isolated using an anti-Tamm-Horsfall protein (THP) antibody, coupled with an anti-rabbit magnetic secondary antibody. They demonstrated positive staining for THP itself and the Na-K-2Cl cotransporter (NKCC2), as verified through immunofluorescent and lectin affinity-fluorescent confocal microscopy techniques. All five lines were immortalised by human terminal transferase (hTERT)-containing lentiviral infection and selected with G418. Both DBA- and THP-isolated DT cells were further distinguished by their absence of staining for markers typical of human proximal tubule cells, such as lotus tetragonolobus lectin (LTA), CD13, and N-cadherin, as well as the collecting duct marker L1CAM.

In 2020, Dr. Tetsuro Kusaba and his team developed a novel technique for isolating hDCT cells from freshly voided urine, leading to the generation of four urine-derived hDCT cell lines (12). In brief, freshly voided urine was centrifuged to obtain cell pellets, which were cultured to allow colony formation. Targeted colonies were selectively isolated using cloning rings. These cells were immortalised through the introduction of SV40T. Western blot analysis confirmed the cellular origin of these urine-derived renal epithelial cells, showing positive expression for the canonical DCT marker, NCC. It also verified the absence of markers indicative of other renal cell types, including Nephrin (a podocyte marker), THP (a marker of thick ascending limb), Megalin (a proximal tubule marker), and Aquaporin-2 (AQP2, a collecting duct marker). This approach not only demonstrates the viability of non-invasively obtaining renal cells but also illuminates the potential of urine-derived cells for advancing our understanding of renal physiology and disease mechanisms. The availability of hDCT cell models is crucial for advancing renal research and understanding the complexities of kidney physiology and pathophysiology. These cell lines offer a valuable platform for dissecting the molecular mechanisms underlying ion transport, hormonal regulation, and the response to pharmacological agents within the distal tubule.

As informative as these models may be in traditional 2D cell culture, it is desirable to replicate the three-dimensional architecture of the kidney tubule to properly model its physiology. This has begun to be addressed by the development and refinement of kidney organoids and organs-on-a-chip, which are more physiologically relevant models than traditional 2D cultures (70–72). Kidney tubular organoids, derived from both pluripotent stem cells and adult kidney cells, have at least partially recapitulated the complex architecture and specialised roles of various tubular segments (73–76). These organoids have facilitated insights into developmental biology, disease modeling, and the potential for regenerative medicine applications. Likewise, the organ-on-a-chip technology has emerged as a powerful tool for simulating the tubular microenvironment with precise control over biological and mechanical conditions. Kidney-on-a-chip models have been instrumental in studying the dynamics of drug transport, nephrotoxicity, and disease mechanisms at an unparalleled resolution (77, 78). They enable the replication of the fluid flow and shear stress conditions inherent to the renal tubules, providing a dynamic understanding of cellular behaviors and interactions in a more *in vivo*-like context (79, 80). Together, these cutting-edge cell culture approaches have significantly expanded current capability to model kidney diseases, test pharmacological interventions, and explore the intricacies of kidney function and pathology, opening new avenues for personalised medicine (81, 82).

### Limitation

This review is an extensive analysis of the available literature on DCT cell models. However, there are several limitations of this review. A significant challenge was the incomplete reporting of methodologies across many articles, particularly concerning culture conditions. This lack of detailed methodology made it difficult to extract comprehensive information and formulate optimal culture recipes. Notably, over half of the articles reviewed did not specify the antibiotics and supplements used, including their concentrations. This omission complicates the determination of whether these components were not used or merely unreported. Thus, the culture conditions summarised in tables may be skewed towards studies with more transparent and thorough protocols.

The issue of incomplete reporting extends to the review of transfection methodologies and their effectiveness. A 78% of papers failed to report the transfection state (transient or stable) and only a handful of papers (6 in total) provided details on the efficiency of transfection. Given these significant gaps in reporting, it is impractical for this review to recommend a specific transfection method for each cell line based on efficiency.

## GRANTS

Keith Siew: This research was funded in whole, or in part by the Wellcome Trust [Grant number 110282/Z/15/Z]. For the purpose of open access, the author has applied a CC BY public copyright licence to any Author Accepted Manuscript version arising from this submission.

Alessandra Grillo: This fellowship was supported by Kidney Research UK [RP_017_20190306].

## AUTHOR CONTRIBUTIONS

Chutong Zhong, Zhen Sun: Conceived and designed research, performed experiments, analysed data, interpreted results of experiments, prepared figures, drafted manuscript

Alessandra Grillo: Edited and revised manuscript

Keith Siew: Conceived and designed research, edited and revised manuscript, approved final version of manuscript.

Stephen B Walsh: Conceived and designed research, edited and revised manuscript, approved final version of manuscript

